# Modulation of ferroptosis sensitivity by TXNRD1 in pancreatic cancer cells

**DOI:** 10.1101/2020.06.25.165647

**Authors:** Luke L. Cai, Richard A. Ruberto, Matthew J. Ryan, John K. Eaton, Stuart L. Schreiber, Vasanthi S. Viswanathan

## Abstract

The selenoprotein thioredoxin reductase 1 (TXNRD1) plays a central role in ameliorating oxidative stress. Inhibition of TXNRD1 has been explored as a means of killing cancer cells that are thought to develop an enhanced reliance on such antioxidant proteins. In the context of ferroptosis, a non-apoptotic form of oxidative cell death, TXNRD1 has been proposed to cooperate with the phospholipid hydroperoxidase enzyme glutathione peroxidase 4 (GPX4) to protect cells from the lethal accumulation of lipid peroxides. Here, we report our unexpected finding that in pancreatic cancer cells, CRISPR–Cas9-mediated loss of TXNRD1 confers protection from ferroptosis induced by small-molecule inhibition of GPX4. Insights stemming from mechanistic interrogation of this phenomenon suggest that loss of TXNRD1 results in increased levels of GPX4 protein, potentially by influencing availability of selenocysteine, a scarce amino acid required by both proteins for proper synthesis and function. Increased abundance of GPX4 protein, in turn, protects cells from the effects of small-molecule GPX4 inhibition. These findings implicate selenoprotein regulation in governing ferroptosis sensitivity. Furthermore, by delineating a relationship between GPX4 and TXNRD1 contrary to that observed in numerous other settings, our discoveries underscore the context-specific nature of ferroptosis circuitry and its modulators.

## Introduction

Ferroptosis is a non-apoptotic, iron-dependent mode of cell death resulting from uncontrolled lipid peroxidation and destruction of polyunsaturated phospholipids.^1–3^ In recent years, ferroptosis induction has garnered considerable therapeutic interest; in particular, as a means through which to selectively kill therapy-resistant cancer cells.^4,5^ Functional genetic approaches to map the cell circuitry underlying ferroptosis vulnerability of such cancer cells have identified core and conserved pathways that control the peroxidation of polyunsaturated fatty acids (PUFAs) as well as their incorporation into phospholipids.^6–9^ However, these same efforts have highlighted the existence of lineage- and cell state-specific regulators of ferroptosis biology that lie outside of the central PUFA pathway.^4,9^ Further characterization of these factors can improve understanding of the ways in which context-specific programs interplay with the biology of lipid peroxidation. These insights can additionally inform the development of patient stratification strategies for clinical translation of ferroptosis-inducing therapies and can help anticipate context-specific mechanisms of resistance to such therapies.

Here, we report on thioredoxin reductase 1 (TXNRD1) as a ferroptosis modulator in pancreatic cancer cells. TXNRD1 is a cytosolic flavin adenine dinucleotide oxidoreductase that relies on an uncommon selenocysteine residue to reduce the active-site disulfide in thioredoxins.^10–13^ This C-terminal selenocysteine operates alongside NADPH and FAD domains to reduce oxidized species.^14^ The unique enzyme structure of TXNRD1, combined with the reactivity of its selenocysteine residue toward smallmolecule electrophiles,^15–18^ warrants careful consideration when dissecting the function of TXNRD1 through genetic versus pharmacological inhibition. Inhibition of TXNRD1 has been previously explored as a therapeutic approach through which to kill cancer cells possessing a heightened dependence on resolving oxidative stress.^19^ In particular, inhibition of TXNRD1 has been proposed to increase sensitivity of cancer cells to other perturbations that contribute to increased oxidative stress.^20^

In this manuscript, we identify and explore the connection between TXNRD1 and the ferroptosis pathway, and uncover a relationship between perturbation of TXNRD1 and sensitivity to ferroptosis that runs contrary to the above paradigm. Specifically, we find that genetic loss of TXNRD1 confers protection against smallmolecule inhibition of glutathione peroxidase 4 (GPX4), potentially via regulatory feedback mechanisms governing selenoprotein biosynthesis. We investigate this interaction within pancreatic cancer cell lines and probe its relevance more broadly in cancer cell lines of this lineage. This unexpected result contributes to a growing understanding surrounding the nuances of ferroptosis biology and emphasizes the importance of context specificity when considering the targets and mechanisms responsible for conferring decreased susceptibility to ferroptosis. In conjunction with general themes that emerge, this particular finding should be borne in mind when developing single-agent and combinatorial strategies for inducing ferroptosis.

## Results and Discussion

### Genome-scale CRISPR screening implicates TXNRD1 in ferroptosis

Our research group has performed genome-scale CRISPR–Cas9 ferroptosis suppressor screens in cancer cell lines of many different lineages including kidney, melanoma, and pancreatic cancer. The goal of these investigations has been to identify genes which, when knocked out, confer resistance to smallmolecule inhibitors of GPX4, the critical regulator of ferroptosis.^2^ Such efforts have reliably uncovered core determinants of PUFA–phospholipid biology, including acyl-CoA synthetase long chain family member 4 (*ACSL4*) and lysophosphatidylcholine acyltransferase 3 (*LPCAT3)* as strong positive regulators of ferroptosis sensitivity.^9,21^ In addition, these screens have highlighted the existence of lineage-specific modulators of ferroptosis sensitivity. The contribution of the hypoxia-inducible factor (*HIF)* pathway in ferroptosis sensitivity of clear-cell renal cell carcinoma and the role of cytochrome P450 oxidoreductase (*POR)* enzyme in ferroptosis sensitivity of melanoma serve as compelling examples of this phenomenon.^9,21^ In the context of pancreatic cancer, a CRISPR–Cas9 ferroptosis suppressor screen using ML210 (Figure 1A) performed in KP-4, a pancreatic ductal adenocarcinoma (PDAC) cell line, identified several candidate lineagespecific regulators (Table 1). The presence of *TXNRD1* in this list attracted our attention because such a finding is at odds with general perspectives within the field on the link between the thioredoxin and ferroptosis pathways. The function of TXNRD1 as an antioxidant protein would suggest a cooperative role in promoting cell survival alongside GPX4. Instead, the result from our CRISPR–Cas9 screen suggests an opposite relationship; namely, that loss of TXNRD1 confers decreased sensitivity to small-molecule GPX4 inhibition.

**Figure 1.**
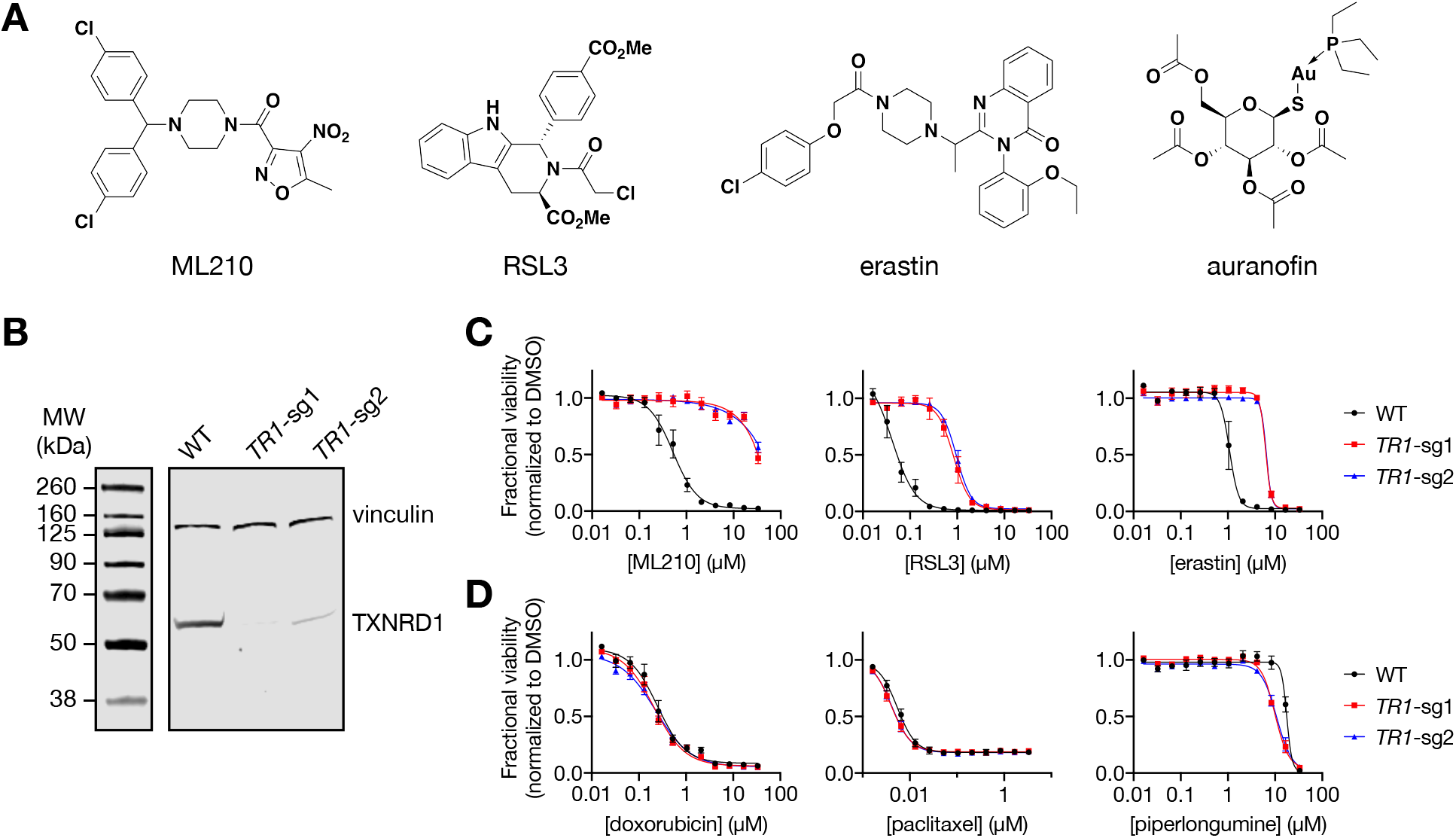
Genetic deletion of TXNRD1 decreases sensitivity to ferroptosis inducers. (A) Chemical structures of ferroptosis inducers (ML210, (1*S*,3*R*)-RSL3, and erastin) and the TXNRD1 inhibitor auranofin. (B) Western blot analysis of TXNRD1 protein levels in wild-type (WT) KP-4 cells and those exposed to RNPs containing sgRNAs targeting *TXNRD1* (*TR1-sg1* and *TR1* -sg2). Vinculin was used as a loading control. Data are representative of two biological replicates. (C) Dose-response curves for WT or RNP-transfected KP-4 cells treated with ferroptosis inducers. Data are plotted as mean ± s.e.m., *n* = 4 technical replicates. (D) Dose-response curves for WT or RNP-transfected KP-4 cells treated with non-ferroptotic cell-killing compounds. Data are plotted as mean ± s.e.m., *n* = 4 technical replicates.

**Table 1.** Top hits identified from CRISPR-Cas9 ferroptosis suppressor screen performed with KP-4 cells.

| gene symbol | median log<sub>2</sub> fold change | median rank percentile |
| --- | --- | --- |
| *ACSL4* | 6.651 | 0.013 |
| *ATP2C1* | 4.610 | 0.222 |
| *LPCAT3* | 4.822 | 0.164 |
| *TMEM164* | 3.122 | 1.492 |
| *TMEM165* | 4.433 | 0.335 |
| *TXNRD1* | 4.476 | 0.241 |

Gene symbols colored in gray were also uncovered as hits in previous screens.

### TXNRD1 knockout confers decreased sensitivity to ferroptosis-inducing small molecules

To validate this result, we first transfected KP-4 cells with Cas9-sgRNA ribonucleoprotein (RNP) complexes containing guide sequences targeting *TXNRD1*. This approach yielded a heterogeneous mixture of cells which, in bulk, displayed drastically reduced TXNRD1 protein levels, as assessed by western blot (Figure 1B). In concurrence with results from the initial genome-scale screen, knockout of TXNRD1 in these cells conferred decreased sensitivity to the ferroptosis-inducing small molecules ML210, (1*S*,3*R*)-RSL3 (hereafter referred to as RSL3), and erastin (Figures 1A,C and S1–S3).^1,2,22^ This decreased sensitivity was specific to ferroptosis inducers, as TXNRD1 loss did not impact cell killing by a range of other compounds representing targeted therapies, general ROS inducers, and other cytotoxic substances (Figures 1D and S1–S3). These results were recapitulated in another PDAC cell line, PANC-1, in which ablation of TXNRD1 yielded similar protection against ferroptosis, thereby suggesting that this phenomenon may be more broadly extensible to other pancreatic cancer cells (Figures S4 and S5).

### Knockout of TXNRD1 does not impact target engagement of GPX4 by covalent inhibitors

To explore the mechanisms through which TXNRD1 loss confers protection from ferroptosis, we first sought to establish whether small-molecule ferroptosis inducers could still bind their respective targets. This line of inquiry was especially important given that the thioredoxin system is known to play a key role in the metabolism of xenobiotic substances.^23^ Such transformations can be critical for activation of prodrug molecules; for example, the covalent GPX4 inhibitor ML210 requires conversion to its active form in live cells.^22^ We assessed direct interaction between GPX4 and ML210 through label-free cellular thermal shift assay (CETSA)^24,25^ within intact KP-4 WT and *TXNRD1*^−/−^ cells. We observed thermal stabilization of GPX4 in the ML210-treated condition irrespective of TXNRD1 knockout (Figure 2A,B). This finding suggests that differential target engagement does not underlie resistance to ML210 in *TXNRD1*^−/−^ cells. Furthermore, these results establish that TXNRD1 is not necessary for unmasking ML210 to yield its active form.

**Figure 2.**
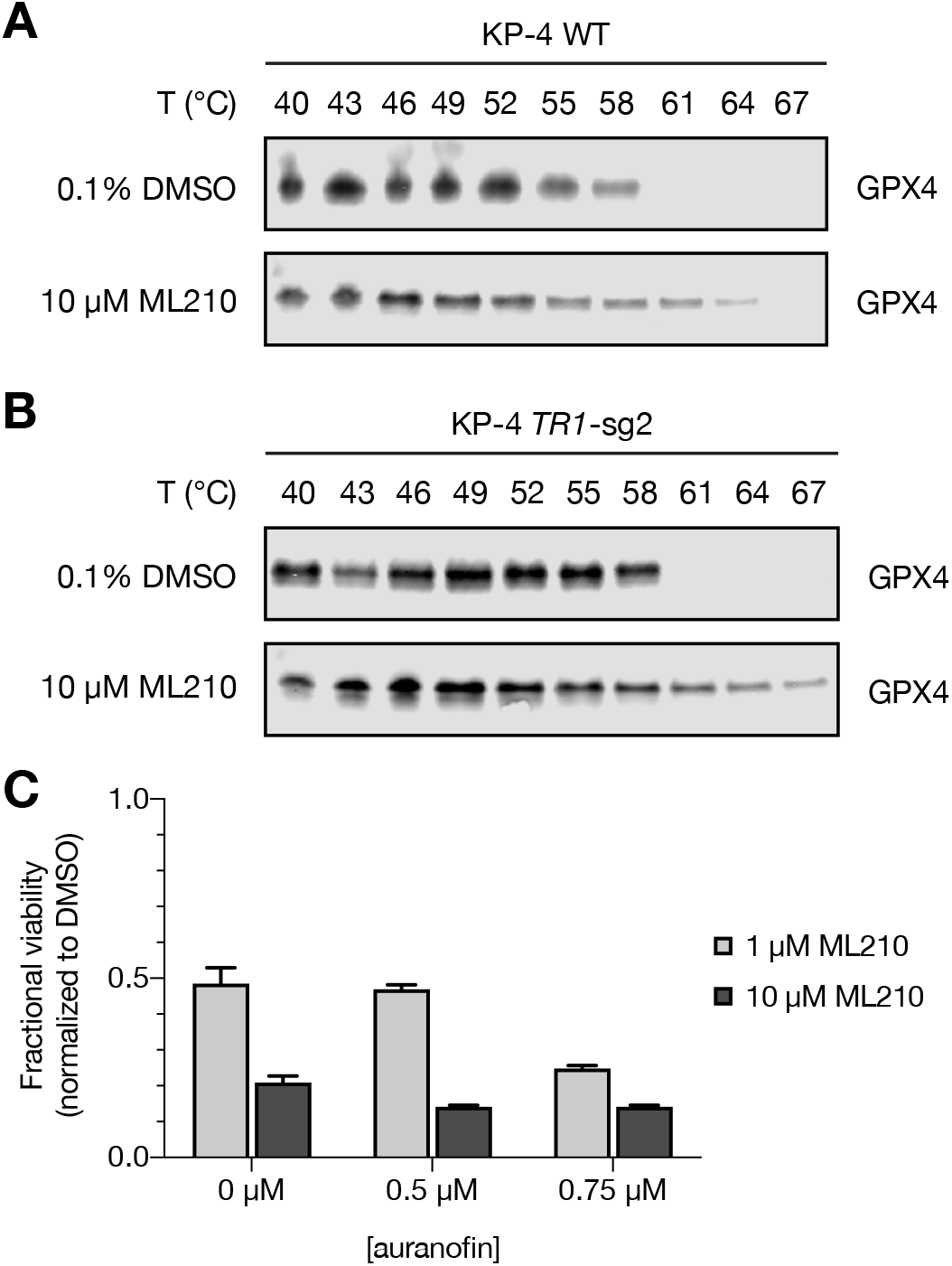
Knockout of TXNRD1 does not block target engagement or cell death induced by covalent GPX4 inhibitors. (A) GPX4 CETSA for intact KP-4 WT cells treated with vehicle or ML210 for 1 h. (B) GPX4 CETSA for intact KP-4 TXNRD1 knockout cells treated with vehicle or ML210 for 1 h. (C) Viability of KP-4 cells co-treated with auranofin and ML210. Data are plotted as mean ± s.d., *n* = 4 technical replicates.

### Inhibition of TXNRD1 does not protect cancer cells from ferroptosis inducers

We next sought to distinguish between two mechanistic possibilities underlying protection of *TXNRD1*^−/−^ cells from ferroptosis inducers: inhibition of TXNRD1 enzymatic function versus decreased abundance of the protein itself, which notably contains selenocysteine. First, we determined whether small-molecule inhibition of TXNRD1 phenocopies genetic loss of the protein with respect to decreased susceptibility to ferroptosis induction. To assess this possibility, we treated KP-4 cells with ML210 and auranofin (Figure 1A), a gold-containing small molecule that targets TXNRD1. The metal center of auranofin binds the selenium atom of the active-site selenocysteine in TXNRD1 and perturbs associated oxidoreductase pathways.^26–28^ Pharmacological inhibition of TXNRD1 with auranofin pre-treatment failed to rescue cell death induced by ML210 (Figure 2C).

### GPX4 protein levels are increased by TXNRD1 knockout

These results encouraged us to investigate whether decreased TXNRD1 protein levels may lead to a compensatory increase in abundance of GPX4 protein through previously reported regulatory feedback mechanisms involving selenocysteine availability. Briefly, selenocysteine biosynthesis occurs cotranslationally and involves the conversion of seryl-tRNA^[Ser]Sec^ to selenocysteinyl-tRNA^[Ser]Sec^ through an ATP-dependent, two-step process.^29,30^ This dependence on selenium availability suggests that the synthesis of one selenoprotein can impact the synthesis of other selenocysteine-containing proteins by affecting total selenium availability. Given that increased abundance of GPX4 is sufficient to confer decreased sensitivity to ferroptosis,^2,31,32^ the expression levels of other selenoproteins have important implications for ferroptosis sensitivity. Indeed, previous reports detail increased GPX4 protein levels in the setting of genetic TXNRD1 perturbation.^33,34^ To investigate this hypothesis, we assessed GPX4 protein levels in *TXNRD1*^−/−^ cells through western blot, and confirmed higher GPX4 protein expression in these cells compared to WT (Figure 3A). We also validated the ability of increased GPX4 protein levels to prevent ferroptotic cell death (Figures 3B, S6, and S7). Importantly, increased GPX4 protein levels did not rescue loss in viability induced by other cytotoxic substances (Figures 3C, S6, and S7), suggesting that this effect is specific to ferroptotic cell death.

**Figure 3.**
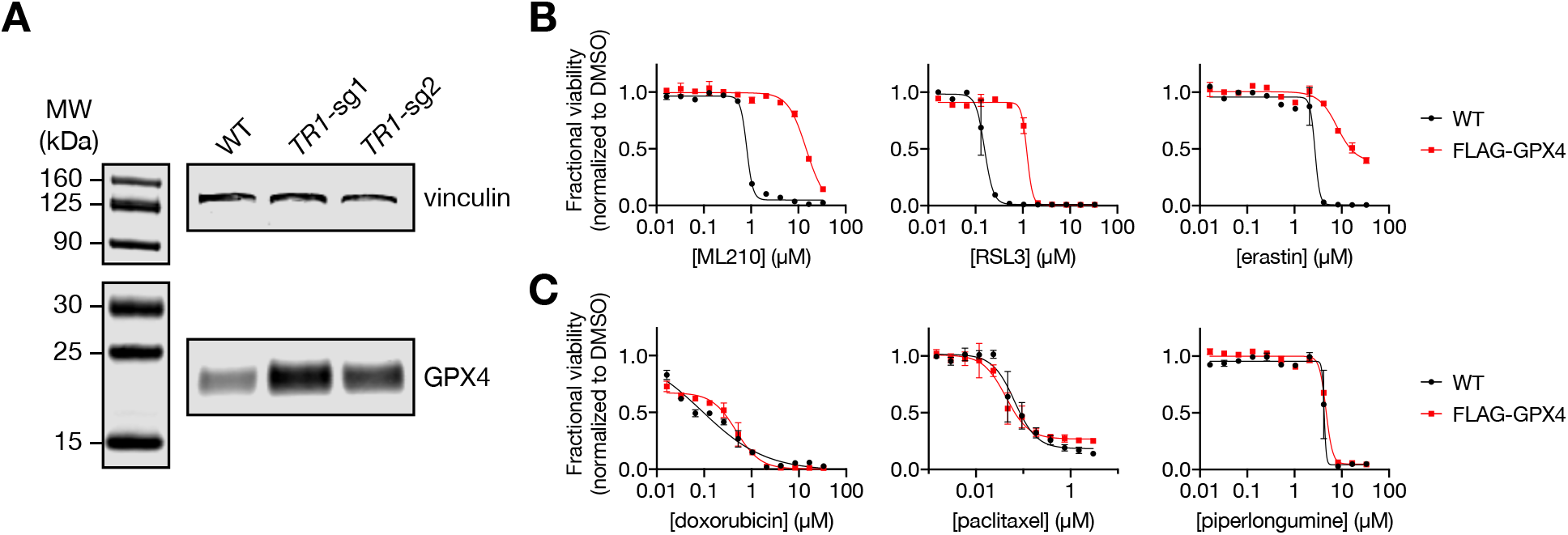
Increased GPX4 protein levels rescue cell-killing effects of ferroptosis inducers. (A) Western blot analysis of GPX4 protein levels in KP-4 WT cells and those exposed to RNPs containing *TXNRD1*-targeting guide RNAs. Vinculin was used as a loading control. Data are representative of two biological replicates. (B) Dose-response curves for WT or FLAG-GPX4 expressing LOX-IMVI cells treated with ferroptosis inducers. Data are plotted as mean ± s.e.m., *n* = 2 technical replicates. (C) Dose-response curves for WT or FLAG-GPX4 expressing LOX-IMVI cells treated with non-ferroptotic cell-killing compounds. Data are plotted as mean ± s.e.m., *n* = 2 technical replicates.

### TXNRD1 loss does not rescue cell death caused by GPX4 knockout

A key prediction of the mechanism described above is that while TXNRD1 loss can confer protection against smallmolecule ferroptosis inducers, such a change would not rescue cells from genetic loss of GPX4 itself. To investigate this hypothesis, we generated KP-4 cells in which both GPX4 and TXNRD1 were knocked out. These cells were initially maintained in growth medium containing the radical-trapping antioxidant ferrostatin-1 (fer-1), which functionally prevents ferroptotic cell death stemming from loss of GPX4.^35,36^ Upon withdrawal of fer-1 from growth medium, all populations of *GPX4^−/−^* cells, including those that were *TXNRD1^−/−^*, exhibited substantial losses in viability (Figure 4). We speculate that the presence of a small number of surviving cells under these specific experimental conditions reflects incomplete knockout of both genes in a heterogeneous, bulk population.

**Figure 4.**
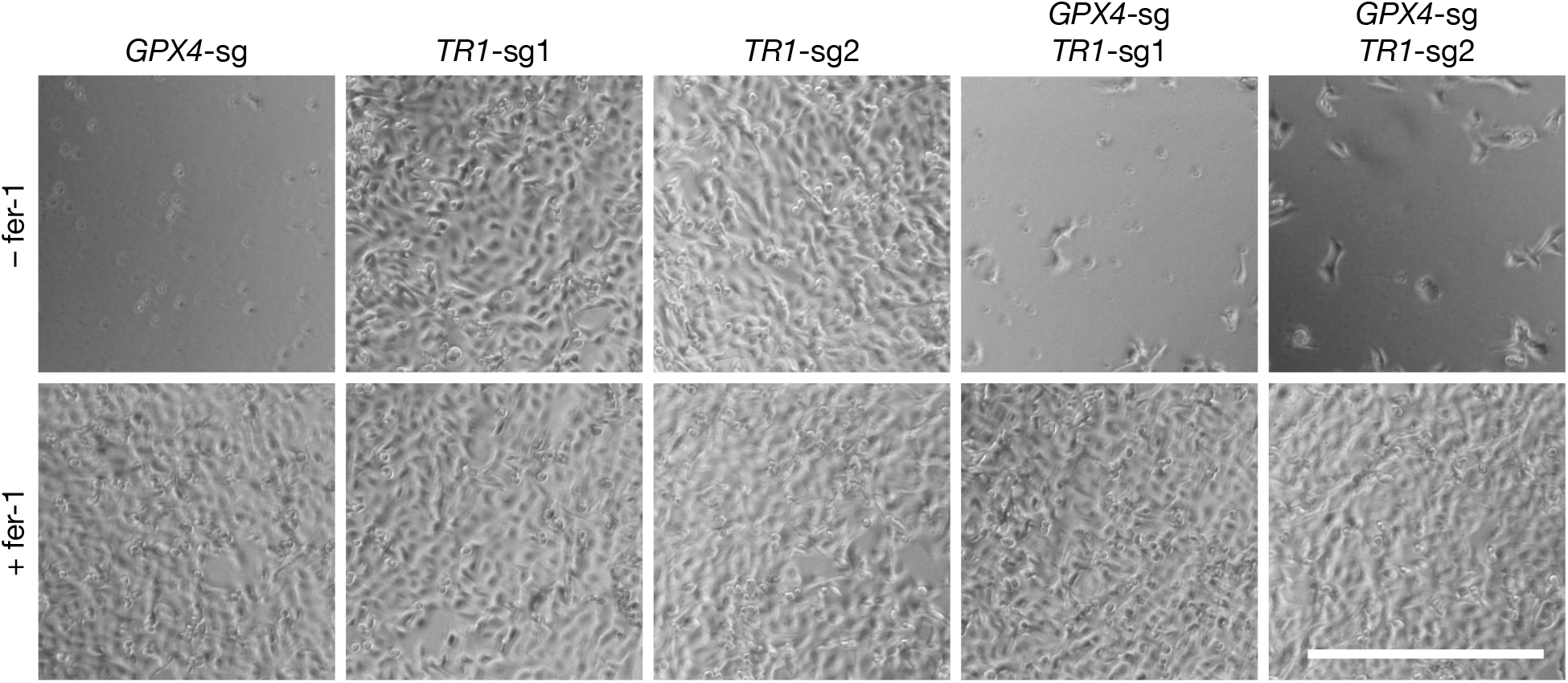
TXNRD1 knockout in the background of GPX4 depletion fails to rescue from cell death. images of *GPX4^−/−^, TXNRD^−/−^*, and *GPX4^−/−^/TXNRD1^−/−^* KP-4 cells 24 h after withdrawal of fer-1 from growth medium (top row) or 24 h after culture in medium containing fer-1 (bottom row). Scale bar, 500 μm.

### TXNRD1 protein levels correlate with ferroptosis sensitivity

Given the ferroptosis protection conferred by TXNRD1 loss in two PDAC cell lines, we speculated that this GPX4–TXNRD1 axis may constitute a mechanism underlying the spectrum of intrinsic ferroptosis sensitivities observed across pancreatic cancer cell lines. Accordingly, correlating transcriptomic data with an area-under-the-curve (AUC) metric of the cell-killing activities of ML210 and RSL3 (Cancer Dependency Map at https://depmap.org/portal/)^37–40^ reveals that pancreatic cancer cell lines expressing lower amounts of TXNRD1 mRNA are less susceptible to cell death induced by GPX4 inhibitors (Figure 5).

**Figure 5.**
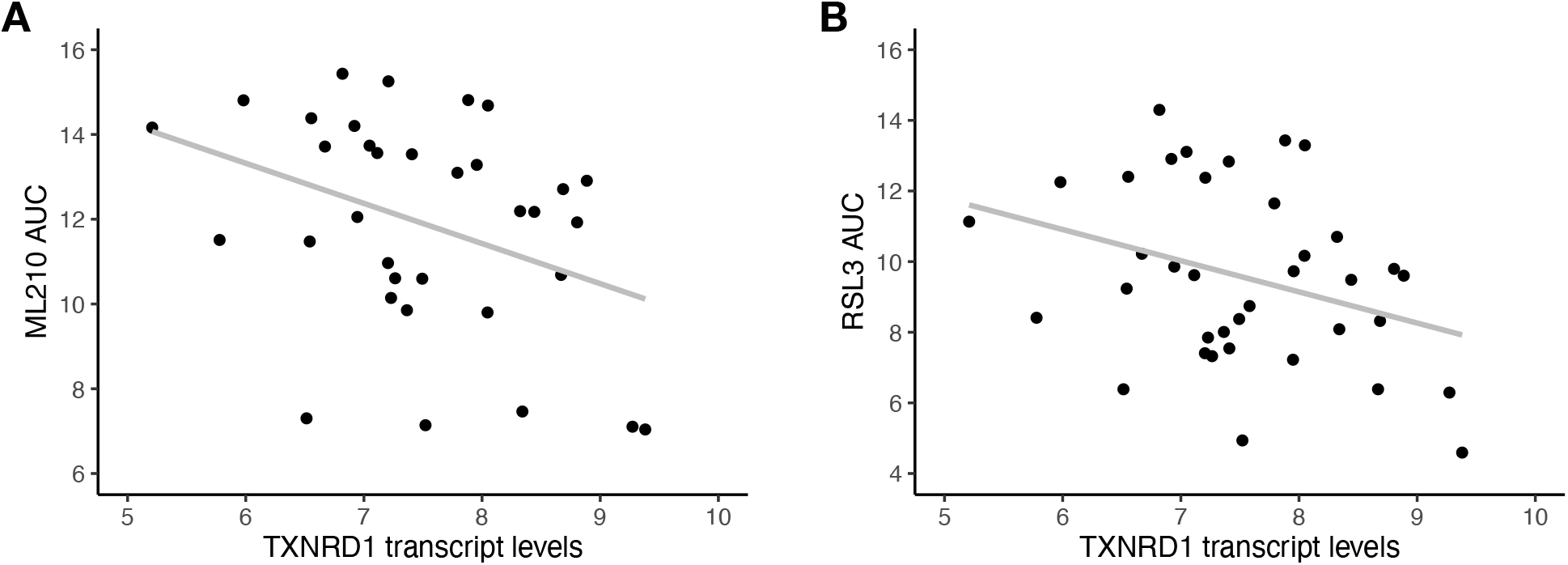
Increased TXNRD1 expression levels are associated with increased sensitivity to covalent GPX4 inhibitors in pancreatic cancer cell lines. (A) Correlation between ML210 AUC and TXNRD1 mRNA levels presented as log_2_(TPM + 1). Lower AUC values correspond to increased sensitivity to ML210. Each point represents a pancreatic cancer cell line. Pearson correlation coefficient *r* = −0.37 with *p* = 0.031. (B) Correlation between RSL3 AUC and TXNRD1 mRNA levels presented as log_2_(TPM + 1). Each point represents a pancreatic cancer cell line. Pearson correlation coefficient *r* = −0.33 with *p* = 0.044.

## Conclusions

Factors regulating sensitivity to ferroptosis are only beginning to be understood, and they show a large degree of context specificity involving lineage and oncogene programs. Here, we identify TXNRD1 as a strong negative modulator of ferroptosis susceptibility in pancreatic cancer cells. To our knowledge, no other genome-scale CRISPR–Cas9 ferroptosis suppressor screen has yielded *TXNRD1* as a significant hit. This result highlights the context-specific nature of ferroptosis modulators and illustrates that even paradoxical relationships are possible within this pathway. Our findings, particularly as they relate to TXNRD1, point toward a novel and largely overlooked mechanism of interaction between ferroptosis and thioredoxin biology, and emphasize the importance of selenoprotein regulation as a key determinant of ferroptosis sensitivity.

## Materials and Methods

### Cell lines

KP-4 and PANC-1 pancreatic as well as LOX-IMVI melanoma cancer cells were obtained from the Broad Institute Biological Samples Platform. KP-4 and PANC-1 cells were maintained in DMEM (Gibco). LOX-IMVI cells were maintained in RPMI 1640 medium (Gibco). All media were supplemented with 10% fetal bovine serum (Gibco), 100 U/mL penicillin (Gibco), and 100 μg/mL streptomycin (Gibco). LOX-IMVI FLAG-GPX4 cells were grown in medium additionally supplemented with 50 nM sodium selenite. Cultures were maintained at 37 °C in a humidified 5% CO_2_ atmosphere. All cells were periodically tested for mycoplasma using the MycoAlert Plus kit (Lonza) and confirmed to be free of contamination.

### Compounds

ML210 and (1*S*,3*R*)-RSL3 were prepared in accordance with previously published procedures.^2,22,41^ Doxorubicin, erastin, paclitaxel, and piperlongumine were purchased from SelleckChem. Auranofin, ferrostatin-1 (fer-1), and sodium selenite were purchased from Sigma-Aldrich. Fer-1 was used at a concentration of 1.5 μM for all experiments.

### Generation of knockout cells

Knockout cells were generated using a ribonucleoprotein (RNP) transfection method.^42^ Briefly, cells were transfected with RNP complexes comprised of EnGen Cas9 NLS, *S. pyogenes* (New England Biolabs); Alt-R CRISPR–Cas9 crRNA; and Alt-R CRISPR– Cas9 tracrRNA (Integrated DNA Technologies) with Lipofectamine RNAiMAX transfection reagent (Invitrogen) according to the manufacturer’s instructions. Target sequences were as follows: CCAGGGATGCCCAAGTAACG, TTACCCCATCTAGTTCCAAG for *TXNRD1*; and CACGCCCGATACGCTGAGTG for *GPX4*. Knockout was confirmed via western blotting and functional assessment.

### Cell viability assays

Viability experiments were performed in 384-well format using opaque, white tissue-culture-treated plates (Corning). Cells were seeded at 1000 cells/well in 30 μL of growth medium and allowed to adhere for 24 h after which they were exposed to compounds using a CyBi-Well vario pin-transfer instrument (Analytik Jena AG). Following a 72-h treatment time, cellular ATP levels, as a surrogate for viability, were quantified using CellTiter-Glo reagent (Promega) and an EnVision plate reader (PerkinElmer). Data were analyzed in R, and doseresponse curves and plots were generated in GraphPad Prism 8 as previously described.^39^

To determine the effects of TXNRD1 inhibition on GPX4 dependence, cells were seeded at 25% confluence in 12-well plates in the presence of auranofin (0, 0.5, or 0.75 μM). These cells were then allowed to adhere overnight before being exposed to a range of ML210 concentrations for 24 h.

To assess the effects of fer-1 withdrawal on GPX4 and TXNRD1 knockout cells, cells were seeded in 12-well plates at 1 × 10^5^ cells/well in 1 mL of growth medium containing fer-1. After 24 h, medium was aspirated and cells were washed with PBS (3 × 1 mL). Growth medium without fer-1 was added to the wells and cells were incubated for 24 h. Cells were visualized with an EVOS FL cell imaging system (Thermo Fisher Scientific).

### Western blotting

Adherent cells were trypsinized, pelleted by centrifugation, washed with PBS, and flash frozen in liquid nitrogen. Cells were then lysed in PBS containing 1% Triton X-100 (Sigma-Aldrich) and protease inhibitor cocktail (Roche) for 20 min on ice with intermittent vortexing. Lysate was centrifuged (20,000 × g, 20 min, 4 °C) to remove insoluble material. Total protein concentration in the supernatant was quantified with a Pierce BCA protein assay kit (Invitrogen). Fifty micrograms of protein were mixed with Laemmli SDS sample buffer (6×, Boston BioProducts) and heated at 95 °C for 10 min. Samples were resolved by SDS-PAGE (Bolt 4– 12% Bis-Tris plus gel, Invitrogen) and transferred to a nitrocellulose membrane using an iBlot 2 gel transfer device (Invitrogen). Membranes were blocked with Odyssey blocking buffer (LI-COR Biosciences) for 1 h, incubated with primary antibodies overnight at 4 °C, and washed with Trisbuffered saline containing 0.1% Tween 20 (TBST, 3 × 5 min). Following incubation with secondary antibodies for 1 h at room temperature and subsequent TBST washes (3 × 5 min), membranes were visualized using an Odyssey CLx imaging system (LI-COR Biosciences). Primary antibodies used: anti-GPX4 (Abcam, ab41787, 1:1000), anti-TXNRD1 (Abcam, ab155790, 1:1000), and anti-vinculin (Abcam, ab130007, 1:1000). Secondary antibodies used: IRDye 680RD donkey anti-mouse (926-68072, LI-COR Biosciences, 1:10,000) and IRDye 800CW donkey anti-rabbit (926-32213, LI-COR Biosciences, 1:10,000).

### Cellular thermal shift assay (CETSA)

Cells were treated with 10 μM ML210 or DMSO control (0.1% v/v) for 1 h at 37 °C. Growth medium was aspirated and cells were washed with PBS (pH 7.4), followed by trypsinization and pelleting by centrifugation (500 × g, 5 min). Cells were then resuspended in PBS, aliquoted into PCR strip tubes (50 μL/condition, 10^6^ cells/condition), and heated in a thermal cycler for 3 min. Afterwards, samples were allowed to cool to room temperature for 3 min. Cell lysis and western blotting were then performed as described above.

## Acknowledgments

We thank Suvruta S. Iruvanti for helpful discussions. This work was supported with funding from the National Cancer Institute’s Cancer Target Discovery and Development (CTD^2^) Network (U01CA217848 awarded to S.L.S.).

## Supplementary Information

**Figure S1.**
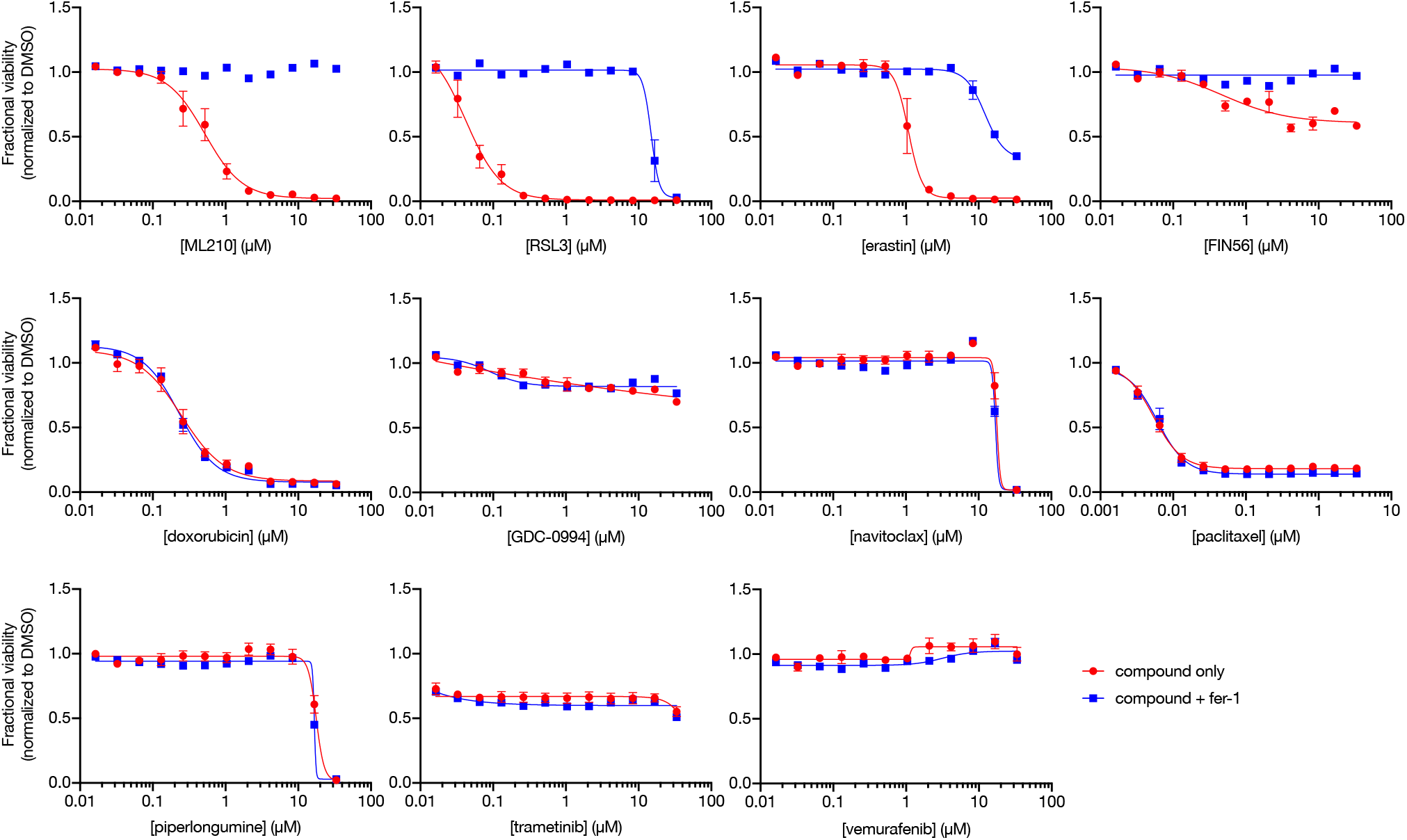
Dose-response curves of KP-4 cells exposed to various cell-killing compounds. Data are plotted as mean ± s.e.m., *n* = 4 technical replicates.

**Figure S2.**
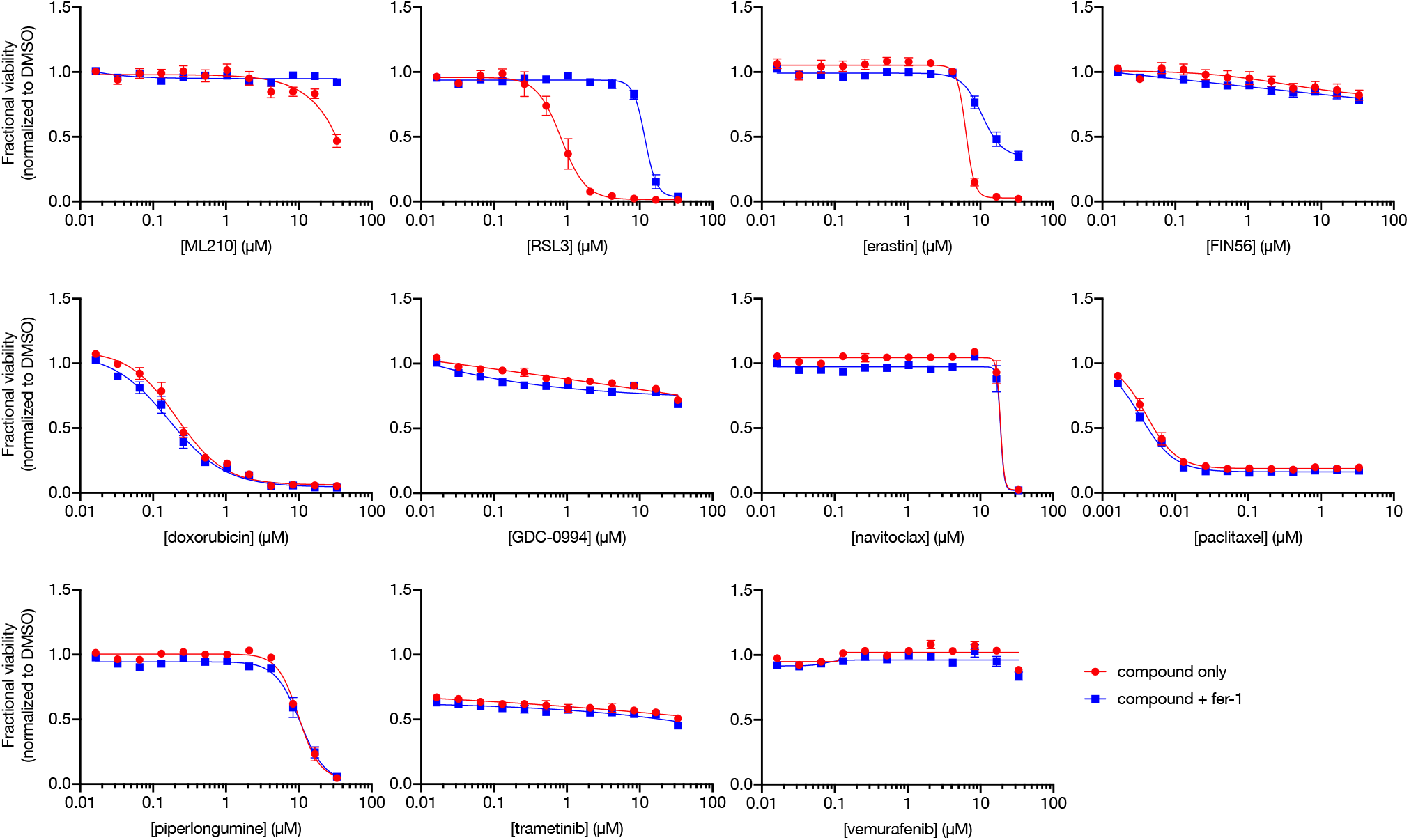
Dose-response curves of KP-4 *TXNRD1*-sgRNA1 cells exposed to various cell-killing compounds. Data are plotted as mean ± s.e.m., *n* = 4 technical replicates.

**Figure S3.**
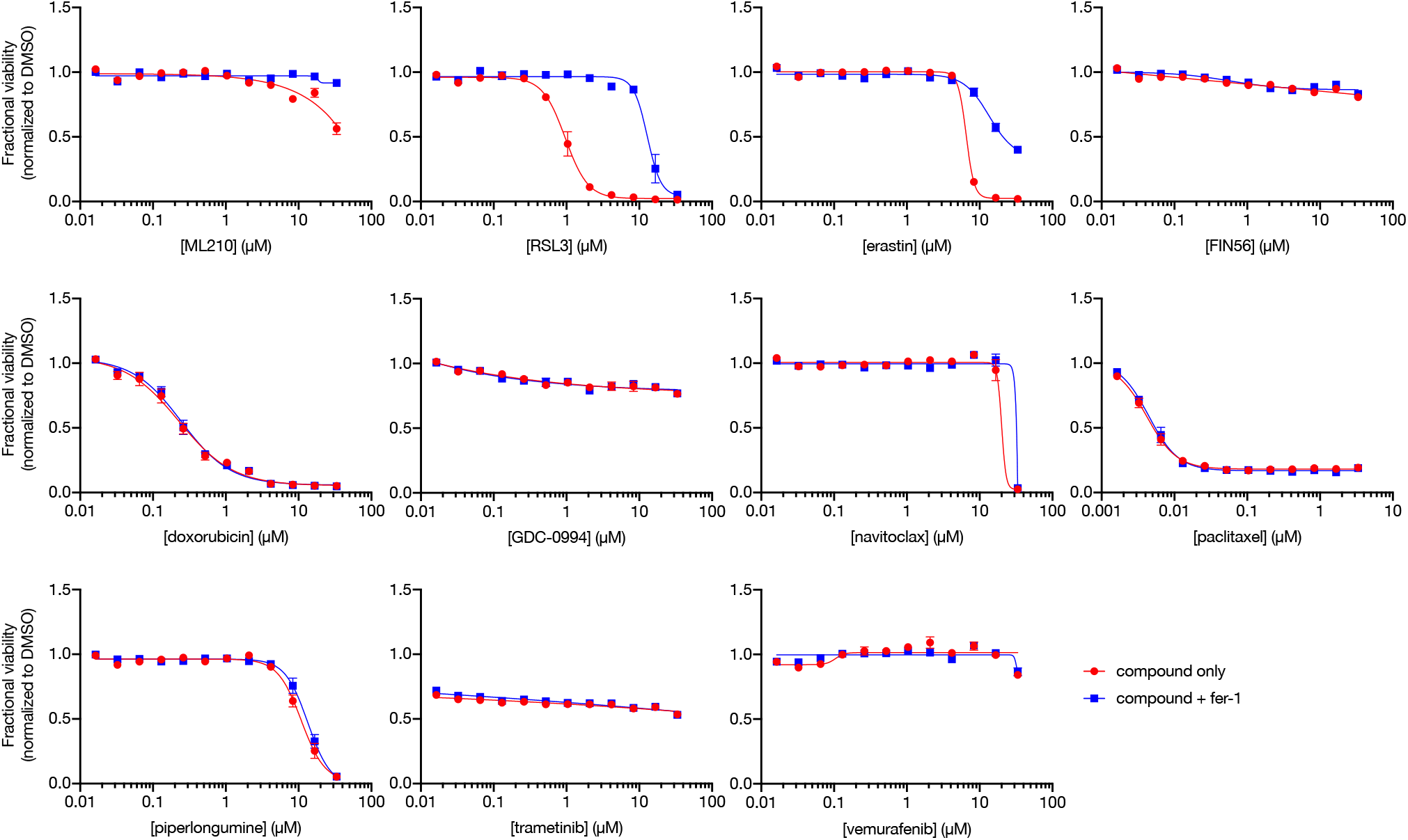
Dose-response curves of KP-4 *TXNRD1* -sgRNA2 cells exposed to various cell-killing compounds. Data are plotted as mean ± s.e.m., *n* = 4 technical replicates.

**Figure S4.**
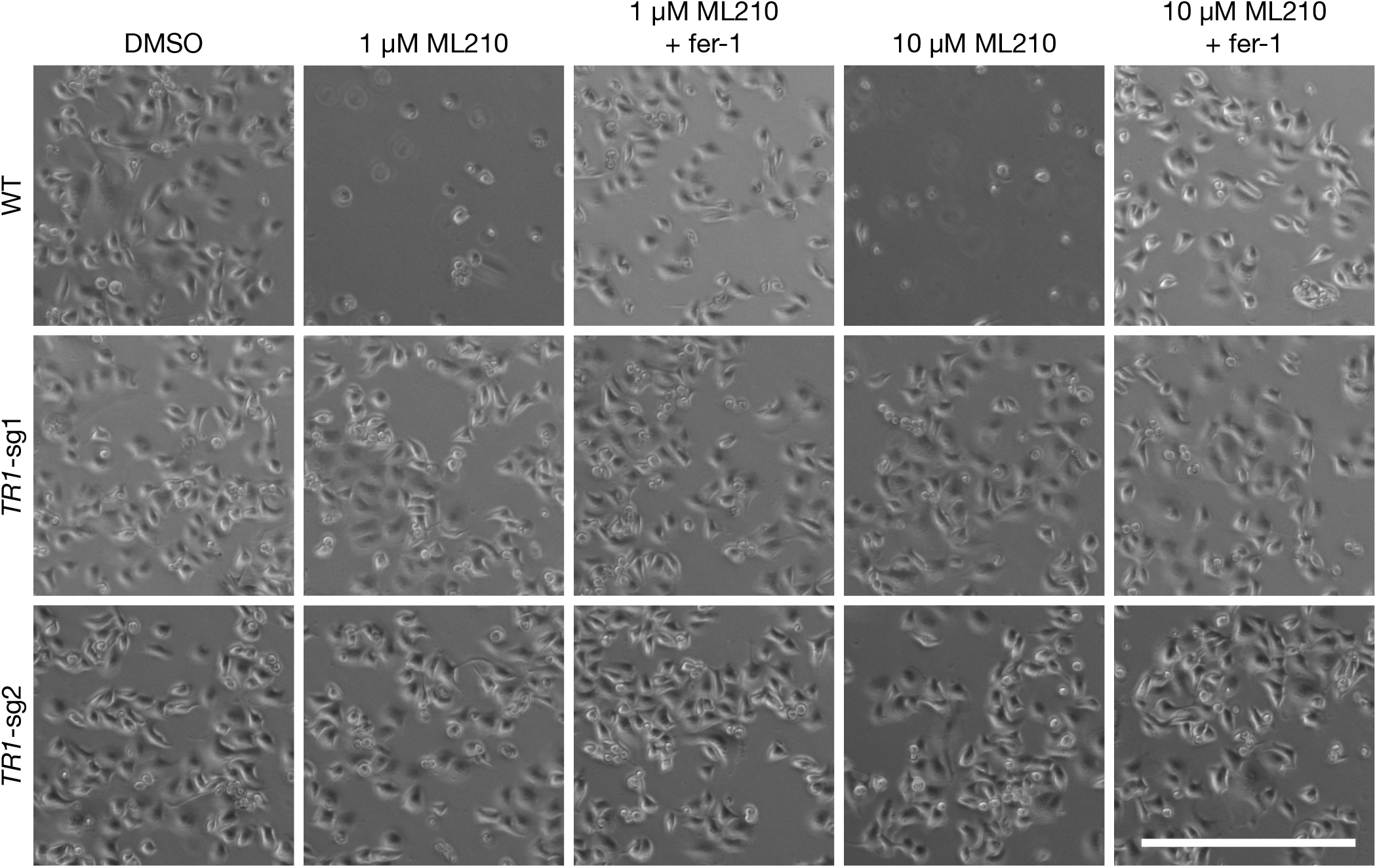
TXNRD1 knockout rescues ML210-induced death in PANC-1 cells. Images of PANC-1 cells after 24 h of exposure to compounds. Scale bar, 500 μm.

**Figure S5.**
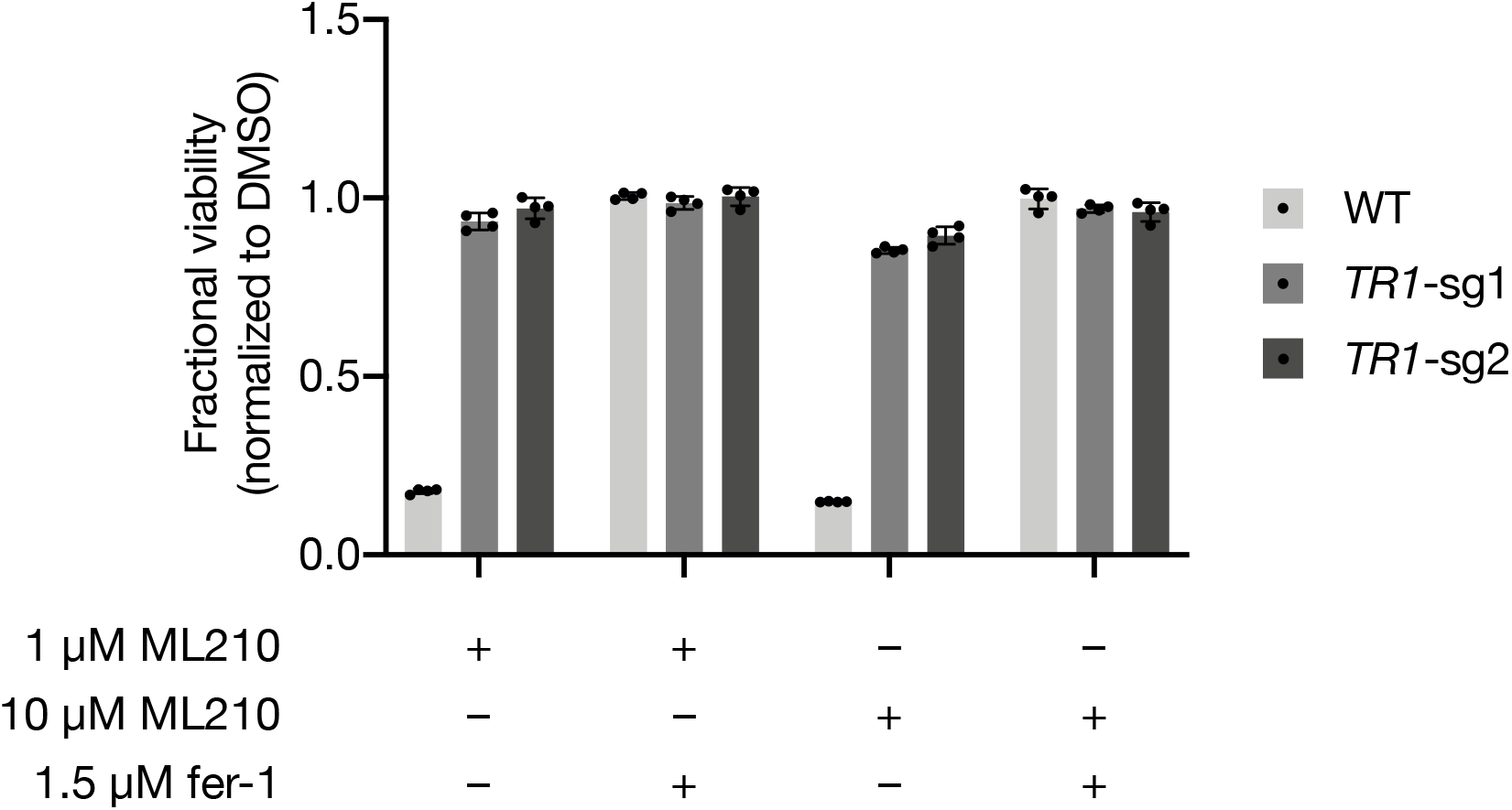
Viability of WT and *TXNRD1^−/−^* PANC-1 cells treated with compounds. Fractional viability data corresponding to images presented in Figure S4. Data are plotted as mean ± s.d., *n* = 4 technical replicates.

**Figure S6.**
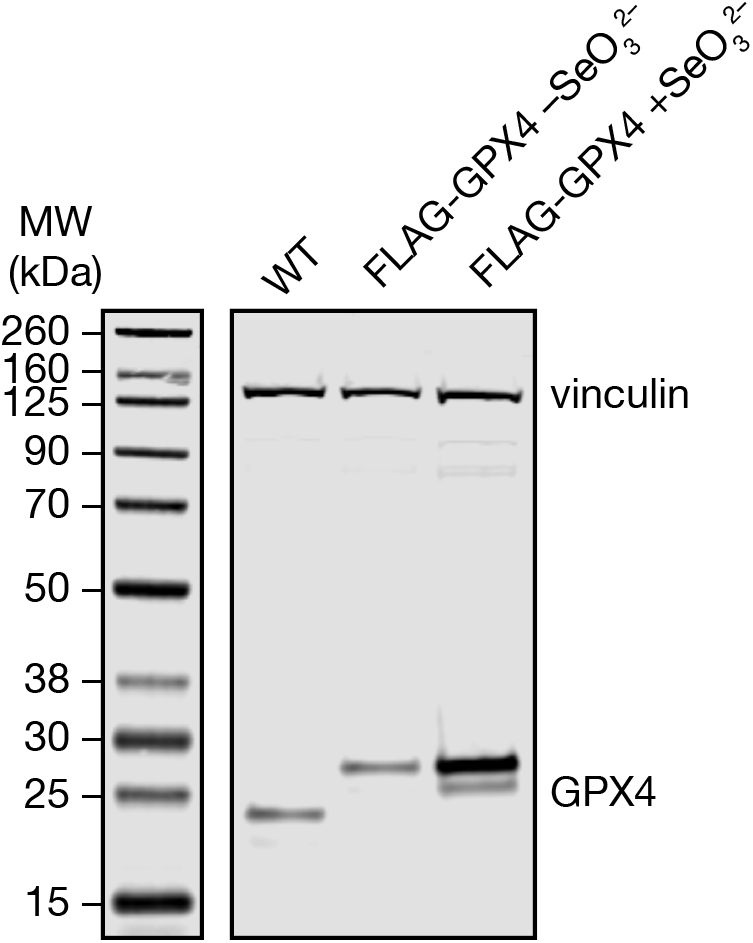
Western blot analysis of GPX4 protein levels in WT and FLAG-GPX4 LOX-IMVI cells. LOX-IMVI cells expressing FLAG-GPX4 constructs were maintained in the absence or presence of 50 nM sodium selenite (Na_2_SeO_3_), as indicated. Vinculin was used as loading control.

**Figure S7.**
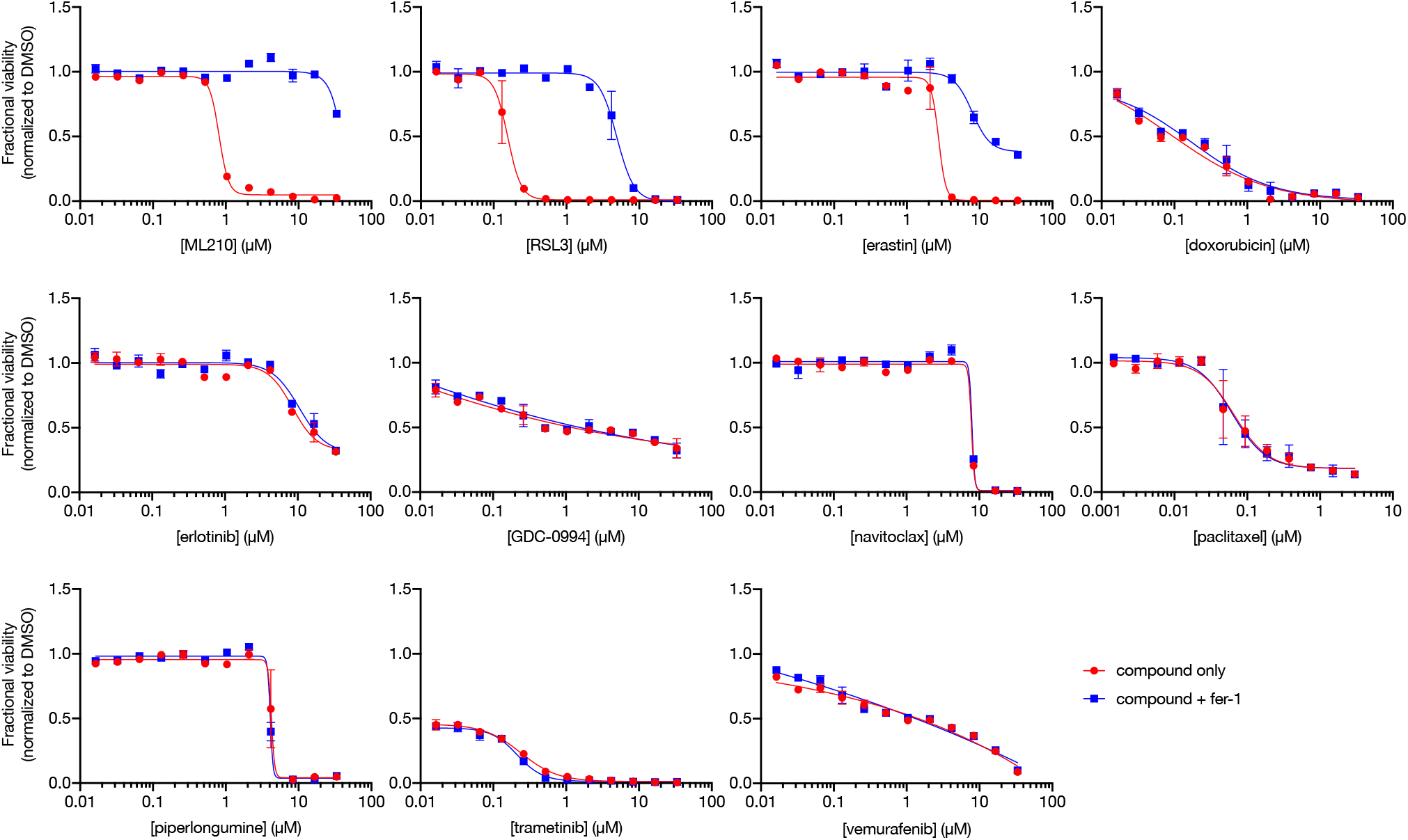
Dose-response curves of LOX-IMVI cells exposed to various cell-killing compounds. Data are plotted as mean ± s.e.m., *n* = 2 technical replicates.

**Figure S8.**
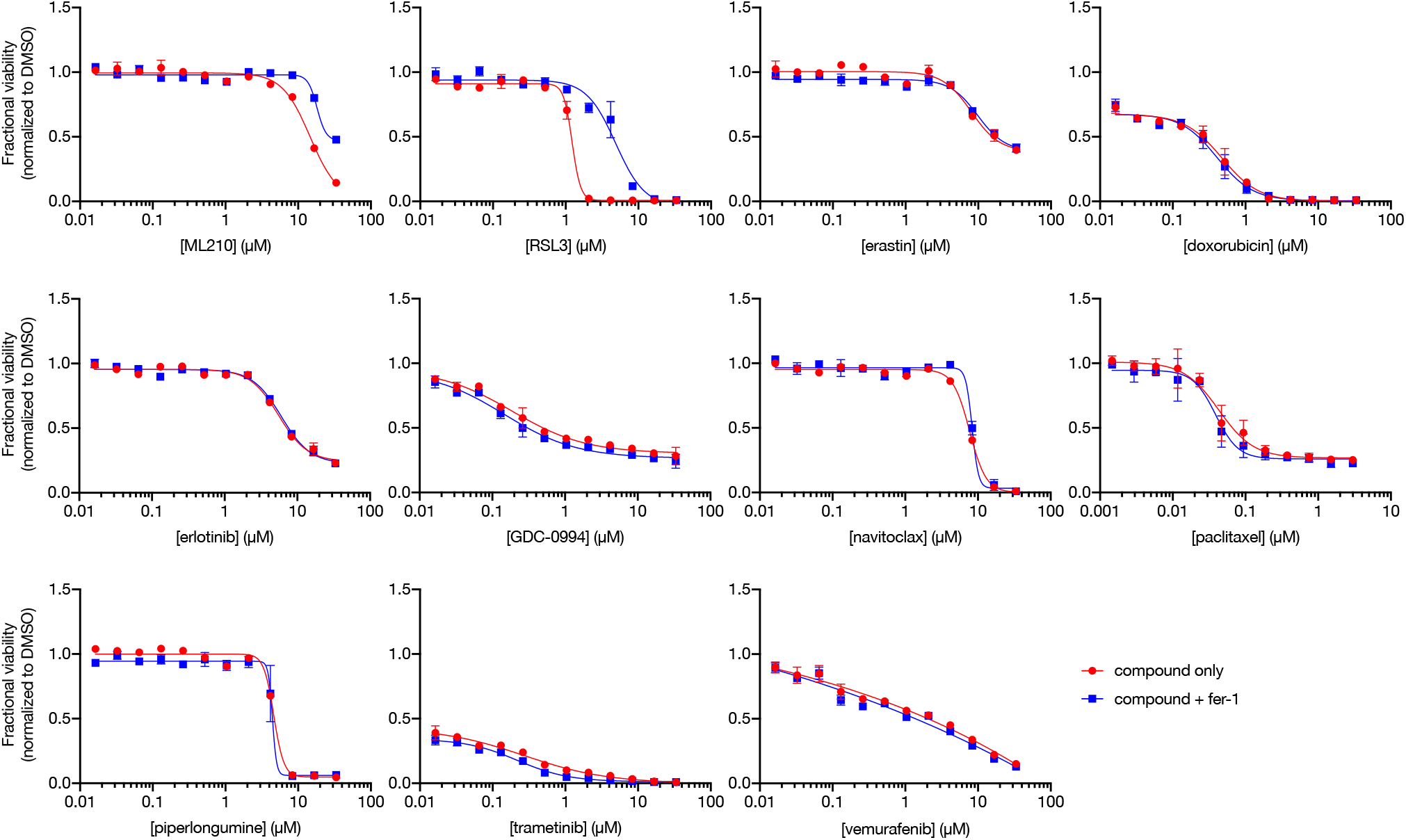
Dose-response curves of LOX-IMVI FLAG-GPX4 cells exposed to various cell-killing compounds. Data are plotted as mean ± s.e.m., *n* = 2 technical replicates.

